# INPP5E regulates CD3ζ enrichment at the immune synapse by phosphoinositide distribution control

**DOI:** 10.1101/2021.01.06.425541

**Authors:** Tzu-Yuan Chiu, Fu-Hua Yang, Weng Man Chong, Hsing-Chen Tsai, Won-Jing Wang, Jung-Chi Liao

## Abstract

The immune synapse, a specialized interface formed between T lymphocytes and antigen-presenting cells (APCs) after antigen recognition, is essential for T cell activation and the adaptive immune response. It has been shown that this interface shares similarities with the primary cilium, a sensory organelle in eukaryotic cells, although roles of ciliary proteins on the immune synapse remain elusive. In this study, we find that inositol polyphosphate-5-phosphatase E (INPP5E), a cilium-enriched protein responsible for regulating phosphoinositide localization, accumulated at the immune synapse during antigen-specific conjugation or antibody capping, and formed a complex with CD3ζ, ZAP-70, and Lck. Silencing INPP5E in T-cells impaired polarized distribution of CD3ζ at the immune synapse, and correlated with a failure of PI(4,5)P_2_ clearance at the center of the synapse. Moreover, INPP5E silencing decreased proximal TCR signaling, including phosphorylation of CD3ζ and ZAP-70, and finally, attenuated IL-2 secretion. Our results suggest that INPP5E is a new player in phosphoinositide manipulation at the synapse, controlling the TCR signaling cascade.

## Introduction

T-cell activation is essential for the cell-mediated adaptive immune response, which requires occupancy of the T-cell receptor (TCR) with a specific antigen peptide/major histocompatibility complex (pMHC) complex on antigen-presenting cells (APCs) (Smith-Garvin et al., 2009). The interaction between the TCR and the MHC complex results in the formation of the immune synapse (Dustin, 2014). Upon conjugation, engagement of TCR and CD28 relocates to the central area of the immune synapse, named the central supramolecular activating complex (cSMAC), allowing the conduction of proximal TCR signaling. Adhesion molecules, such as αLβ2 (LFA-1) and α4β1 (VLA-4) integrin, move toward the peripheral SMAC (pSMAC) and strengthen the binding affinity to ICAM-1 and VCAM-1 (Bertoni et al., 2018). Finally, F-actin spreads over the APC surface at the external ring of the immune synapse, called the distal SMAC (dSMAC). The proximal TCR signaling requires Src family kinase Lck, which phosphorylates the immunoreceptor tyrosine-based activation motifs (ITAMs) in the ɛ and ζ chains of CD3 (Reth, 1989). These motifs are docking sites for the tyrosine kinase ZAP-70 (zeta-chain-associated protein kinase 70). The recruitment and activation of these molecules trigger the downstream signaling phosphorylation and polarization of proteins, including the linker of activated T cells (LAT) and phospholipase Cγ1 (PLCγ1). Moreover, the formation of the immune synapse promotes polarization of the microtubule-organizing center (MTOC) accompanied by the MTOC-associated organelles to the immune synapse (Lowin-Kropf et al., 1998; Martín-Cófreces et al., 2008). These processes finally prompt the mature immune synapse formation and the functional effector responses, such as secretion of cytokines and exosomes (Huse et al., 2006; Mittelbrunn et al., 2011).

Translocation of MTOC, polarization of MTOC-associated organelles, and concentrated signal transduction activities at the immune synapse prompted researchers to consider their similarity to the primary cilium activities. Griffiths and colleagues refer to the immune synapse as a frustrated cilium(Griffiths et al., 2010). The primary cilium is a protruded structure mediating cellular sensation and essential signaling, transducing molecular activities of sonic hedgehog and Notch signaling, whereas the immune synapse is a flat and circular interface. Interestingly, both structures show microtubule and actin reorganization and centrosome polarization to promote cellular events. For instance, distal centriolar protein CEP164 is polarized and docked to vesicles, facilitating ciliogenesis and ARL13B recruitment in the primary cilium (Schmidt et al., 2012; Stinchcombe et al., 2015). CEP164, ARL13B, and PDE6δ all interact with the inositol polyphosphate-5-phosphatase E (INPP5E), an enzyme essential for maintaining phosphoinositide levels, responsible for INPP5E ciliary targeting (Humbert et al., 2012). Moreover, the interaction between INPP5E and PDE6δ and the cargo release by ARL3, for which ARL13B acts as a guanine nucleotide exchange factor, are necessary for INPP5E enrichment in the primary cilium (Fansa et al., 2016). Notably, ARL3 and ARL13B are also involved in activities of the immune synapse. The membrane-bound Lck is extracted by UNC119A and released at the plasma membrane under the control of the ciliary ARL3 and ARL13B (Stephen et al., 2018). It is unclear whether ARL3 and ARL13B are responsible for certain similarities between the primary cilium and the immune synapse, and if INPP5E may play a role in the immune synapse, such as immune synapse formation and promotion of T cell activation.

Here, we first investigated whether any additional ciliary proteins localized at the immune synapse. We found that several ciliary proteins, including INPP5E, were recruited to the immune synapse. Strikingly, we found that INPP5E interacted with CD3ζ, ZAP-70, and Lck during T-cell activation, controlled CD3ζ recruitment toward the immune synapse, and sustained proximal TCR signaling and IL-2 secretion. The effect of INPP5E on immune synapse formation occurred with PI(4,5)P_2_ redistribution at the immune synapse, suggesting the potential phosphatase role of INPP5E on immune synapse formation. Our data revealed how INPP5E was co-opted as a regulator of early signaling and receptor recruitment during T-cell activation.

## Results

### A group of ciliary proteins are present in T cells

We first tested whether any additional cilium-associated proteins localized close to the immune synapse in T cells. Proteins known to localize at the centriole distal end, the transition zone and the cilium were immunostained in conjugates of Jurkat T-Raji B cells. Localization changes of these proteins were studied upon T cell activation by means of staphylococcal enterotoxin E (SEE) engagement. As expected, without SEE, CD3ζ localized in the cytosol presumably at the regions of the Golgi apparatus/endosomes, whereas with SEE, CD3ζ was recruited to the immune synapse. Centriole protein centrin 2 (CETN), ciliary suppression protein CEP97, and distal appendage protein CEP164 were localized at centrioles, which are close to CD3ζ both in the absence and presence of SEE (Fig 1A). In the presence of SEE, these proteins were polarized to the immune synapse (Lowin-Kropf et al., 1998; Stinchcombe et al., 2015). We also examined cilium specific transition zone proteins, including RPGRIP1L and CEP290 (Garcia-Gonzalo et al., 2011) (Fig 1B). Interestingly, RPGRIP1L and CEP290 localized at the centriole, where they were polarized to the cell contact site during antigen-specific conjugation. That is, at least some of the transition zone proteins were present in Jurkat cells, implying possible involvement of the ciliary machinery in Jurkat T cells.

**Figure 1.**
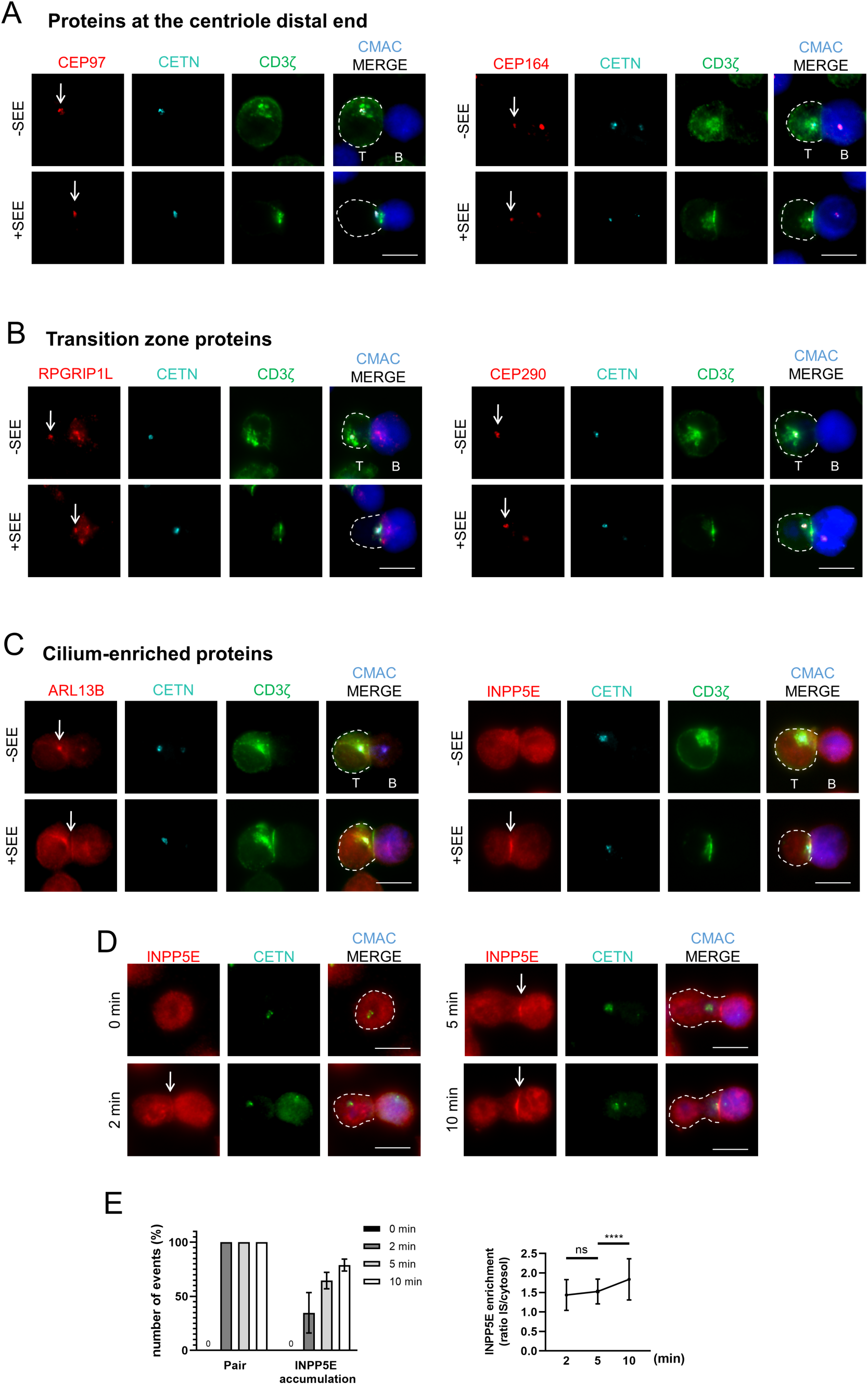
**Localization of cilium-associated proteins in resting and activated T cells.** A-C Immunostaining of (A) proteins at the centriole distal end (CEP97, CEP164), (B) transition zone proteins (RPGRIP1L, CEP290), and (C) cilium-enriched proteins (ARL13B, INPP5E) in conjugates of Jurkat T cells (Left) and CMAC-labeled Raji B cells (Right), in the presence (+) or absence (–) of SEE. Cells were co-stained with anti-centrin 2 (CETN, cyan) and anti-CD3ζ (green) antibodies as centriole and immune synapse markers. Arrows indicate the localization of each cilium-associated protein. Images are representative of at least three independent experiments. D Immunostaining of INPP5E in T cell-APC conjugates with different time points. E Quantification of INPP5E recruitment events (*left* panel) in conjugates (Pair) and INPP5E enrichment ratio (*right* panel) was from two independent experiments. n = 34 for 2 min, n = 70 for 5 min, and n=63 for 10min. 0 min indicated cells were fixed before adding APCs. Scale bar = 10 μm. Error bars indicate mean ± SD. Unpaired student T-test. ns, not significant. ****P < 0.0001.

We further examined cilium-enriched proteins, including ARL13B and INPP5E, to see whether they were adjacent to the immune synapse. We found that ARL13B accumulated at the immune synapse when conjugated with SEE-loaded APCs, similar to the result reported by others (Stephen et al., 2018) (Fig 1C). For the phosphatase INPP5E, we found it localized with the centriole in the absence of SEE, similar to those found in ciliated cells (Xu et al., 2016) (Fig S1A). We also found INPP5E was centered around MTOC at the interphase, and was separated into two parts at the metaphase (Fig S1B). Surprisingly, INPP5E formed a line pattern similar to that of CD3ζ at the interface of T cells and APCs when Jurkat cells were conjugated with SEE-loaded APCs. While INPP5E has been considered as a Golgi-localized enzyme (Gawden-Bone et al., 2018; Kong et al., 2000), the line pattern in Jurkat cells seems to be associated with the cell membrane (Fig S1C). INPP5E also accumulated at the T cell-APC contact site in a time-dependent manner, as INPP5E faintly appeared 2 min after conjugation and continued to accumulate 5-10 min after conjugation (Fig 1D and E). It is thus important to examine whether this cilium-specific phosphatase is indeed associated with the immune synapse.

### INPP5E accumulates at the immune synapse in activated T cells

To validate the INPP5E localization at the immune synapse, we first knocked down endogenous INPP5E by transfecting Jurkat cells with either control or INPP5E-specific siRNA (KD). A clear depletion of INPP5E signals at about 70-kDa was observed in KD cells (Fig 2A). In resting Jurkat cells, enrichment of INPP5E at the centrioles was also depleted in INPP5E-KD cells (Fig S1D). Under SEE treatment, the polarization of INPP5E toward the immune synapse was significantly reduced in INPP5E-KD cells (number of events in conjugation 38.09%, n = 111 versus control cells 71.52%, n = 122) (Fig 2B and C). We then examined INPP5E polarization upon TCR engagement on a planar surface. Cells were first activated with anti-CD3/CD28-coated coverslips for 10 mins, and the recruitment of INPP5E was visualized by total internal reflection fluorescence microscope (TIRF) microscopy and the mean fluorescence intensity (MFI) inside the cells was quantified (Bouchet et al., 2017). Accumulation of INPP5E at the pseudo-synapse was significantly decreased in INPP5E-KD cells when Jurkat cells were activated (Fig 2D and E). We also transfected INPP5E with an N-terminal Flag tag into Jurkat cells and examined the Flag-tagged signals at the T cell-APC contact site. When stimulating with SEE-pulsed APCs, Flag-tagged INPP5E, similar to endogenous INPP5E, was mostly recruited at the synapse (Fig 2F and G), assuring the synapse localization of INPP5E.

**Figure 2.**
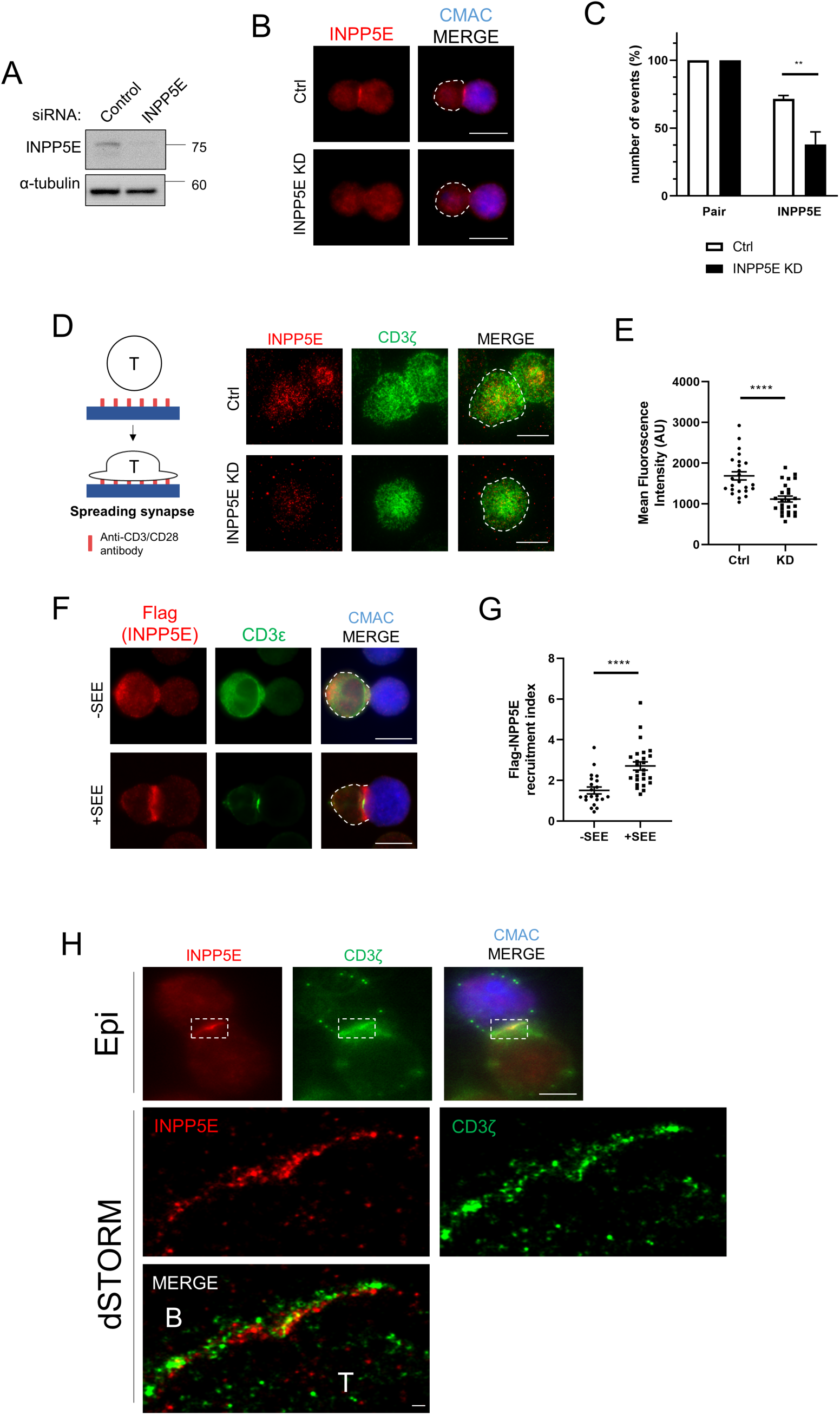
**INPP5E accumulates at the immune synapse in activated T cells.** A-C Jurkat cells were transfected with either control (Ctrl) or INPP5E-specific siRNA (KD). INPP5E expression was analyzed by immunoblotting (A) and immunostaining (B,D). (B) Immunostaining of INPP5E in conjugates of Jurkat T cells and CMAC-labeled SEE-pulsed APCs. Quantification of INPP5E recruitment events (C) in conjugates (Pair) was from four independent experiments. n = 122 conjugates for control, and n = 111 conjugates for INPP5E-knockdown. Scale bar = 10 μm. D-E Jurkat cells were transfected with either control (Ctrl) or INPP5E-specific siRNA (KD). (D) (Left) Schematic figure of the spreading assay. (Right) TIRF imaging of INPP5E localization in spreading Jurkat T cells on anti-CD3/CD28-coated coverslips for 10 min, and measurement of mean fluorescence intensity (E) was from three independent experiments. N = 24 cells for control, and n = 25 cells for INPP5E-knockdown. Scale bar = 5 μm. Error bars indicate mean ± SD. Unpaired student T-test. ns, not significant. **P ≤ 0.0005. ****P < 0.0001. Images are representative of three experiments. F Immunostaining of overexpressed Flag-INPP5E in conjugates of Jurkat T cells and CMAC-labeled SEE-pulsed APCs. Cells were co-stained with anti-CD3ε antibody as the immune synapse marker, and quantification of INPP5E recruitment events (G) in conjugates was from four independent experiments. n = 21 conjugates for -SEE, and n = 26 conjugates for +SEE. Scale bar = 10 μm. H Single molecule images of INPP5E in conjugates of Jurkat T cells and CMAC-labeled SEE-pulsed APCs. Cells were co-stained with anti-CD3ζ antibody as the immune synapse marker. Images are representative of 10 conjugates. Scale bar = 10 μm for epifluorescence (Epi), 200 nm for dSTORM images.

Since the distribution of INPP5E signals mostly appeared at the T cell-APC contact site, it was necessary to examine whether this INPP5E belonged to T cells or B cells. To clarify, we performed super-resolution imaging using direct stochastic optical reconstruction microscopy (dSTORM) to visualize nanoscale localization details at the immune synapse. We found that INPP5E was consistently accumulated at the inner side of Jurkat cells, as the CD3ζ represented the location of the plasma membrane of T cells (Fig 2H). Signals of INPP5E and CD3ζ overlapped in some regions (shown in yellow in the merged image), suggesting possible interactions between them.

### The proline-rich domain is responsible for INPP5E recruitment at the immune synapse

To further map the minimal domain required for INPP5E enrichment at the synapse, we reconstituted different truncations of INPP5E with an N-terminal Flag tag, which has been described in previous studies (Humbert et al., 2012; Plotnikova et al., 2015), into Jurkat cells and observed their localization upon T cell-APC conjugation. We found that the N-terminal portions of INPP5E (amino acids 1-289, 1-604, and 1-626) showed synaptic localization. However, the C-terminal INPP5E (amino acids 79–644 and 288–644) showed impaired synaptic recruitment (Fig 3A, B and C). These results showed that the synaptic recruitment domain of INPP5E was around the N-terminal region, i.e., the proline-rich domain, instead of the ciliary targeting sequence (CTS) of INPP5E known for primary cilia between the inositol polyphosphate phosphatase catalytic domain (IPPc) and the CaaX motif (Fig 3D). Taken together, our results suggested that the proline-rich domain is necessary for synaptic recruitment of INPP5E.

**Figure 3.**
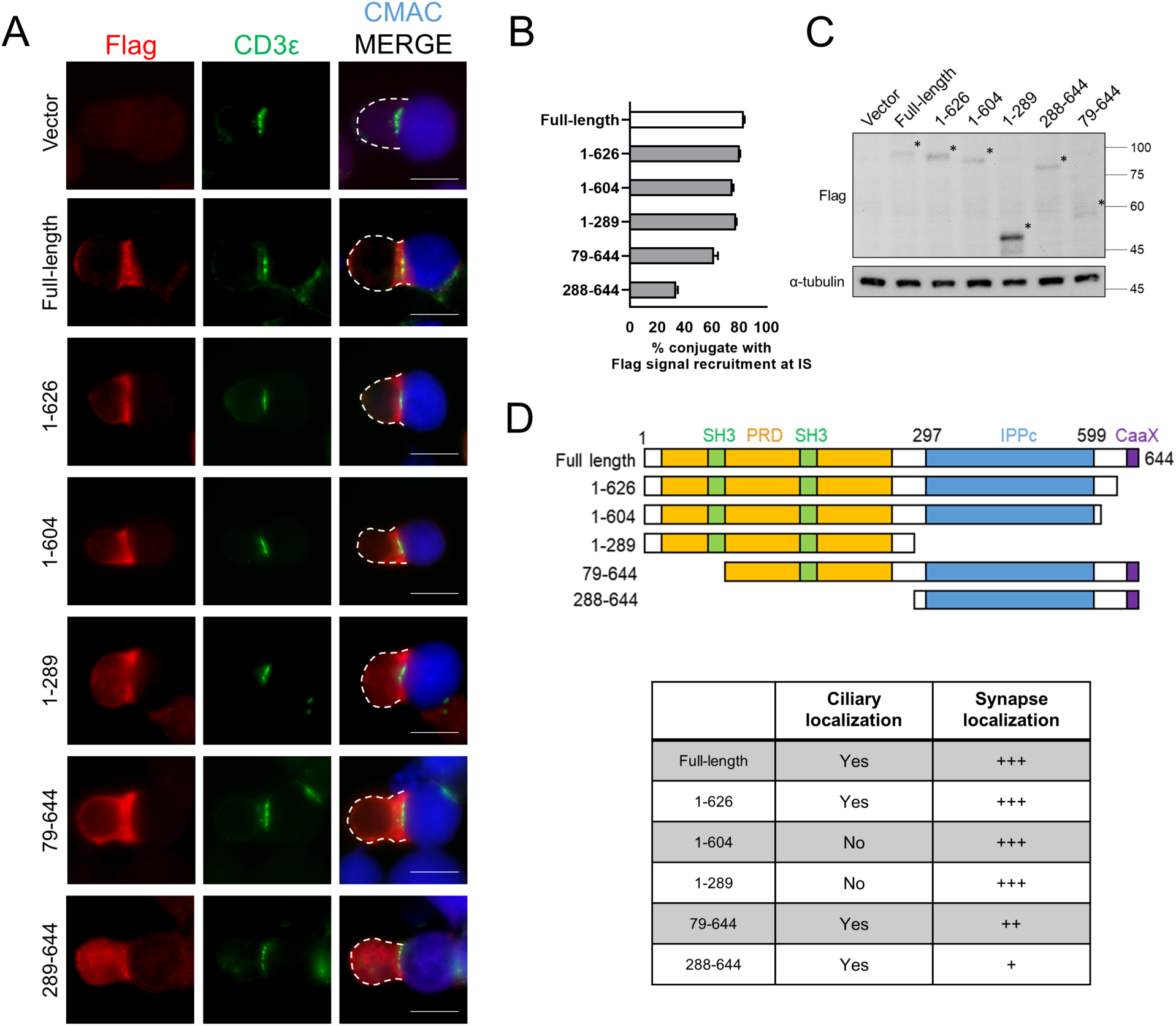
**Proline-rich domain is responsible for INPP5E recruitment at the immune synapse.** A-C Jurkat cells were transfected with expression vectors encoding Flag vector, Flag-INPP5E, and indicated truncations. (A) Immunostaining of Flag-INPP5E truncations in conjugates of Jurkat T cells and CMAC-labeled SEE-pulsed APCs. Cells were co-stained with anti-CD3ε as the immune synapse. Scale bar = 10 μm. (B) Quantification of the Flag-INPP5E truncations recruitment percentage at the immune synapse. n = 30-50 conjugates for each variant. Images are representative of two experiments. (C) Immunoblotting of Flag-INPP5E truncation constructs in Jurkat T cells by 10% SDS-PAGE. The asterisks indicated the transfected proteins. α-tubulin was blotted as internal control. D Summary of INPP5E truncations in primary cilia (Humbert et al., 2012) and immune synapse localization. Proline-rich domain (PRD), SRC Homology 3 Domain (SH3), inositol polyphosphate phosphatase catalytic (IPPc) domain, CaaX motif are marked in orange, green, blue and purple, respectively. The truncations localizing to primary cilia or the immune synapse are labeled on the right. +++, mostly recruitment; ++, medium recruitment; +, low recruitment.

### INPP5E is required for CD3ζ recruitment to the immune synapse

The TCR complex is recruited when stimulating T cells either through APCs pulsed with peptide antigens or anti-CD3/CD28 treatment (Abraham and Weiss, 2004; Lee et al., 2009). The clustering of the TCR/CD3 complex triggers activities of kinases and scaffold proteins to maintain the proximal TCR signaling (Chae et al., 2010; Houtman et al., 2005; Sjölin-Goodfellow et al., 2015) and further to control T cell activation (Minguet et al., 2007). As INPP5E was concentrated at the immune synapse during T cell activation, we examined whether INPP5E played a role in proximal TCR signaling. We capped the TCR/CD3 complex using anti-CD3 and anti-CD28 monoclonal antibodies, a procedure leading to TCR/CD3 clustering (Wülfing et al., 2002). Strikingly, INPP5E interacted with the receptor complex subunit CD3ζ, ZAP-70, and Lck in response to TCR engagement in a time-dependent manner (Fig 4A) (Holdorf et al., 2002; Minguet et al., 2007; Tanimura et al., 2003). To test whether INPP5E could directly interact with these proteins, we used a heterologous HEK293T cell system to approach this question. We first transfected CD3ζ-GFP, Lck, ZAP-70-Myc, and Flag-INPP5E together into HEK293T cells since full phosphorylation of CD3ζ requires Lck and ZAP-70 co-expression in COS-7 or HEK293T cells (James and Vale, 2012; van Oers et al., 2000). CD3ζ-GFP and ZAP-70-Myc were indeed phosphorylated when co-transfecting Lck in HEK293T cells (Fig S2A). Surprisingly, we found that immunoprecipitation of INPP5E could pull down CD3ζ, Lck, and ZAP-70 (Fig 4B). We further showed that INPP5E could interact with these proteins individually by co-expressing CD3ζ, Lck, or ZAP-70 with INPP5E in HEK293T cells. These results suggested that INPP5E could directly interact with these proximal TCR signaling proteins, and the phosphorylation of CD3ζ and ZAP-70 might not be required for their interaction with INPP5E. Notably, the portion of INPP5E that is responsible for interacting with CD3ζ was the C-terminal region (amino acids 288-644). Immunoprecipitation of N-terminal INPP5E (amino acids 1-289) could not pull down CD3ζ-GFP, indicating that INPP5E might utilize different domains for immune synapse docking and CD3ζ interaction (Fig S2B).

**Figure 4.**
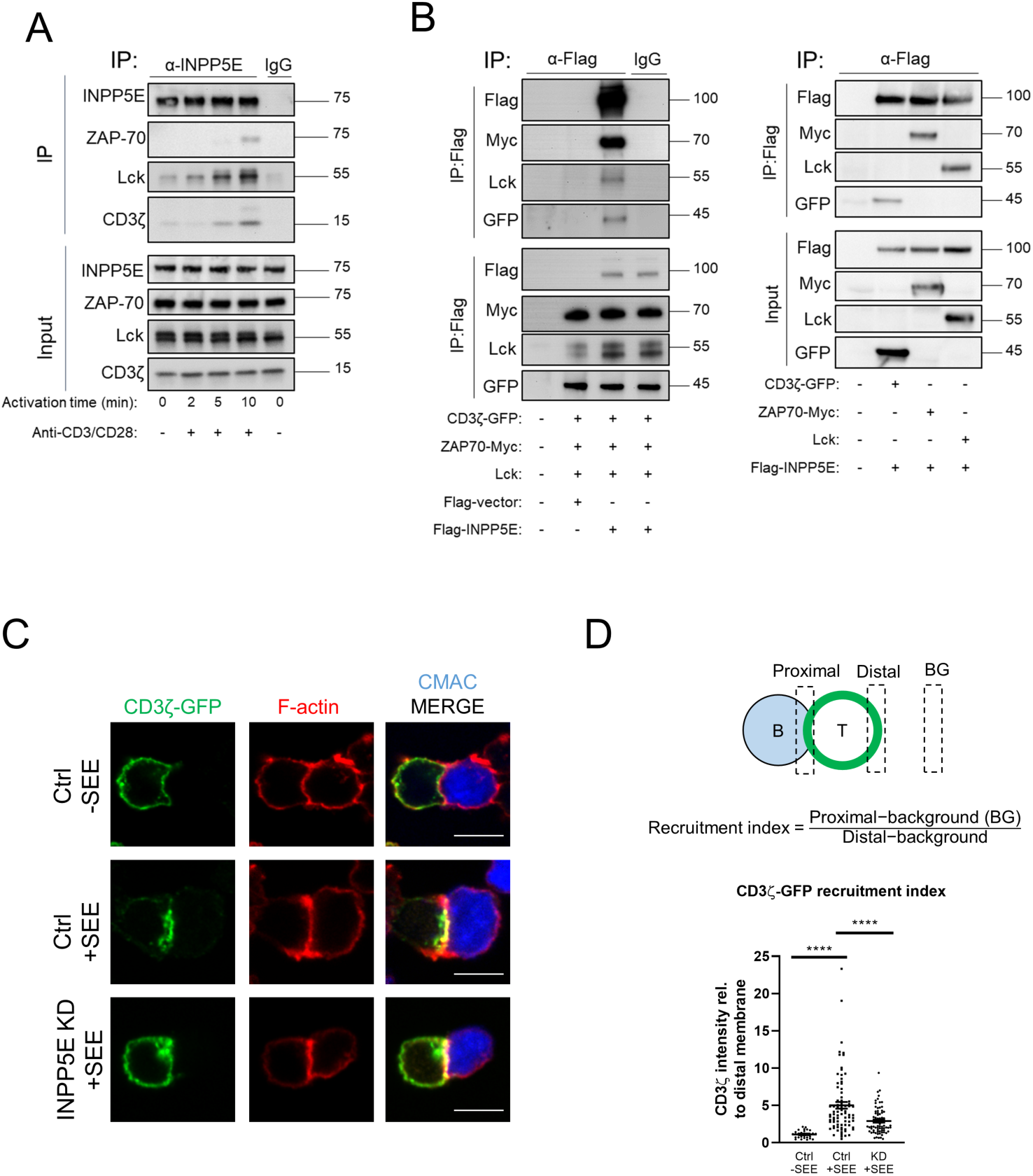
**INPP5E is required for CD3ζ recruitment to the immune synapse.** A Jurkat T cells were activated with crosslinked anti-CD3/CD28 for 2-10 mins, and total cell lysates from each time point were immunoprecipitated (IP) with anti-INPP5E antibody. The immuno-precipitates were resolved by 4-12% SDS-PAGE and immunoblotted with indicated antibodies. IgG lane was used for non-specific antibody control. Images are representative of three experiments. B HEK293T cells were co-transfected with indicated plasmids. Cells were then lysed and immunoprecipitated (IP) with anti-Flag antibody. The immuno-precipitates were resolved by 10% SDS-PAGE and immunoblotted with indicated antibodies. Images are representative of three experiments. C-D Jurkat cells were transfected with either control (Ctrl) or INPP5E-specific siRNA (KD). (C) Immunostaining of CD3ζ-GFP in conjugates of Jurkat T cells and CMAC-labeled APCs, in the absence (–, Upper) or presence (+, middle) of SEE. Cells were co-stained with Alexa Fluor 647 phalloidin to visualize F-actin. Upper and middle, control siRNA transfected cells; lower, INPP5E-specific siRNA transfected cells. Images are representative of four experiments. (D) (Upper) The calculation equation for the relative recruitment index of CD3ζ-GFP to the immune synapse. Dotted black squares indicate the regions that were selected for GFP fluorescence intensity analysis. (Lower) Quantification of CD3ζ-GFP recruitment index in conjugates from four independent experiments. n = 30 conjugates for control (Ctrl) -SEE, n = 81 conjugates for control (Ctrl) –SEE, and n = 73 conjugates for INPP5E-knockdown (KD). Error bars indicate mean ± SD. Unpaired t-test. ****P < 0.0001. Scale bar = 10 μm.

To understand the possible role of INPP5E in immune synapse formation, we reconstituted a CD3ζ construct fused with EGFP into control and INPP5E-KD cells to monitor the exogenous CD3ζ distribution (Blanchard et al., 2002; Soares et al., 2013). We then co-cultured Jurkat cells with APCs before fixation, and the recruitment index of GFP signals at T cell-APC contact sites were analyzed through confocal microscopy (Li et al., 2017). We counted the conjugates with depletion of filamentous actin (F-actin) at the center of the synapse. In the absence of SEE, GFP signals, which were similar to the endogenous CD3ζ localization, were evenly distributed on the plasma membrane (Fig 4C, upper). Interestingly, CD3ζ-GFP was mostly accumulated at the immune synapse in control cells, while the clustering of GFP signals was considerably impaired in the INPP5E-KD cells (recruitment index 2.17, n = 41 versus control cells 4.53, n = 44) (Fig 4C, middle and lower; 4D). Collectively, these data suggested that INPP5E was required for exogenous CD3ζ clustering at the immune synapse.

### INPP5E modulates PI(4,5)P2 distribution to regulate CD3 distribution at the immune synapse

One factor known to regulate TCR/CD3 recruitment is phosphoinositide (Chouaki-Benmansour et al., 2018; DeFord-Watts et al., 2011, 2009). In primary cilia, INPP5E hydrolyzes PI(4,5)P_2_ to PI(4)P to regulate the localization of ciliary Hedgehog signaling inhibitors such as Gpr161 and Tulp3 (Chávez et al., 2015; Conduit et al., 2012; Ukhanov et al., n.d.). Dysregulation of phosphoinositide in INPP5E-deficient mice results in the lethality of embryos and also ciliopathies in humans (Conduit et al., 2012; Jacoby et al., 2009). To test whether INPP5E controls local levels of its substrate at the immune synapse, the sensor expressing EGFP fused to the PI(4,5)P_2_-binding domain of PLCδ (PH-PLCδ) was expressed in control and INPP5E-KD cells (Balla and Várnai, 2009; Szentpetery et al., 2009). We then allowed T-B conjugates to form before fixation, and the 3D structure at the synapse was reconstituted from stacked confocal microscopy images. Cells treated with control siRNA showed negligible PH-PLCδ (Fig 5A and B) fluorescence at the center of the T cell-APC site, indicative of limited PI(4,5)P_2_ molecules at the inner part of the immune synapse. In contrast, the distributions of PH-PLCδ were relatively uniform across the entire contact interface of the immune synapse in the INPP5E-KD cells. That is, the clearance of PIP_3_ and PI(4,5)P_2_ at the inner part of the immune synapse was prevented in INPP5E-KD cells, suggesting that INPP5E controls the PIP_3_ and PI(4,5)P_2_ conversion at the synapse. We also confirmed this observation by transfecting Tubby-GFP, another PI(4,5)P_2_ sensor (Gawden-Bone et al., 2018; Ritter et al., 2017). In control cells, Tubby-GFP signals were again abolished at the synapse; on the other hand, PIP_2_ clearance was impaired in the INPP5E-KD cells, suggesting that INPP5E regulated the PIP_2_ level at the synapse site (Fig S3A and B).

**Figure 5.**
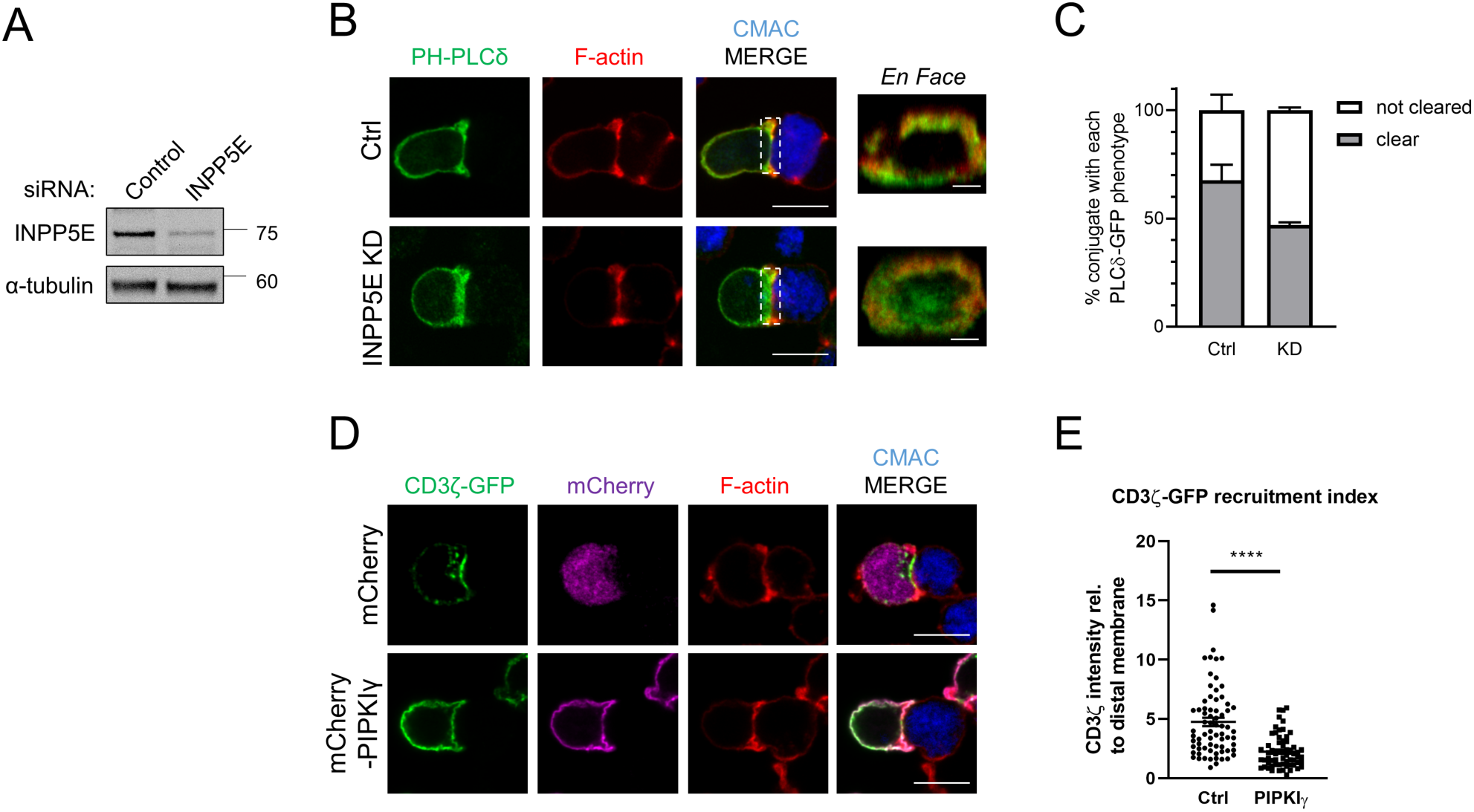
**INPP5E modulates PI(4,5)P*_2_* environment at the immune synapse.** A-C Jurkat cells were transfected with either control (Ctrl) or INPP5E-specific siRNA (KD). (A) INPP5E expression was analyzed by immunoblotting. (B) Immunostaining of PH-PLCδ-GFP in conjugates of Jurkat T cells and CMAC-labeled SEE-pulsed APCs. Images show in the xy plane (scale bar, 10 μm) or 1.0 μm 3D-reconstructions en face at the T cell-APC contact site in the xz plane (scale bar, 3 μm). Cells were stained with the anti-GFP antibody. Alexa Fluor 647 phalloidin was stained to visualize F-actin. Dotted white squares indicate the regions that were selected for 3D reconstruction. Upper, control siRNA transfected cells; lower, INPP5E-specific siRNA transfected cells. Images are representative of three experiments. (C) Quantitation of PH-PLCδ-GFP clearance at the immune synapse in conjugates. n = 54 conjugates for control, and n = 74 conjugates for INPP5E-knockdown cells. D-E Jurkat cells were transfected with expression vectors encoding mCherry and mCherry-PIPKIγ. (D) Immunostaining of CD3ζ-GFP in conjugates of Jurkat T cells and CMAC-labeled APCs, in the presence of SEE. Cells were co-stained with Alexa Fluor 647 phalloidin to visualize F-actin. Upper, mCherry vector transfected cells; lower, mCherry-PIPKIγ transfected cells. Images are representative of three experiments. (E) Quantification of CD3ζ-GFP recruitment index in conjugates from three independent experiments. n = 71 conjugates for mCherry vector transfected cells, n = 62 conjugates for mCherry-PIPKIγ transfected cells. Error bars indicate mean ± SD. Unpaired t-test. ****P < 0.0001. Scale bar = 10 μm.

To assess more directly whether modulating PIP_2_ distribution affected CD3ζ-GFP recruitment, we used phosphatidylinositol-4-phosphate 5-Kinase Type 1 Gamma (PIPKIγ) fused to mCherry to increase PIP_2_ from the plasma membrane (Gawden-Bone et al., 2018). mCherry-PIPKIγ was first introduced into parental Jurkat cells, and cells were then conjugated with SEE-loaded APCs. We found that signals of mCherry-PIPKIγ were evenly distributed at the synapse, which resembled PI(4,5)P_2_ levels (Fig S3A and B). Next, we co-transfected mCherry-PIPKIγ and CD3ζ-GFP into Jurkat cells to visualize CD3ζ distribution under PIP_2_ ablation. Excitingly, recruitment of CD3ζ-GFP was largely diminished in the PIPKIγ-expressed cells, while the clustering of GFP signals remained accumulated at the immune synapse in mCherry control cells (Fig 5C and D). This result indicated that the dysfunction of immune synapse response in INPP5E-KD cells could be caused by negative regulation of PI(4,5)P_2_, similar to the findings in primary cilia (Xu et al., 2016). It is noteworthy that although the clearance of PI(4,5)P_2_ was inhibited when INPP5E was deficient, we found that the localization of F-actin was not visibly affected, different from the outcome when inhibiting a PI(4,5)P_2_ lipase, PLCγ1 (Fig S3C and D) (Gawden-Bone et al., 2018; Ritter et al., 2017). Taken together, these data suggested that INPP5E modulated PI(4,5)P_2_ level at the synapse, and thus affected CD3ζ recruitment.

### INPP5E is essential for proximal TCR signaling and effector functions

TCR recruitment toward the immune synapse is an outcome of T cell activation (Compeer et al., 2018; Vivar et al., 2016). Moreover, dephosphorylation of PI(4,5)P_2_ by exogenous phosphatase Inp54p in mouse T-cell lymphoma enhances the lateral mobility of the TCR/CD3 complex on the plasma membrane, thus increasing TCR activation and signaling (Chouaki-Benmansour et al., 2018). To validate whether INPP5E affects proximal TCR signaling, control and INPP5E-KD Jurkat cells were activated with anti-CD3/CD28 for various durations to stimulate the TCR response (Carpier et al., 2018; Giardino Torchia et al., 2018). At 5 and 15 min, INPP5E-KD cells showed a considerable attenuation of both CD3ζ and ZAP-70 phosphorylation levels upon T cell activation as compared to the response in control cells, while the total amounts of CD3ζ and ZAP-70 were not affected (Fig 6A and B). PLCγ1 phosphorylation was also slightly decreased, though it was statistically insignificant. Moreover, the phosphorylation of CD3ζ and ZAP-70 did not appear to be affected by Lck, as the level of active Lck (Lck Y394) remained similar between control and INPP5E-KD cells. Furthermore, control and INPP5E-KD cells were conjugated with SEE-loaded B cells for 10 mins, and cells were fixed and stained with phospho-CD3ζ and phospho-ZAP-70 antibodies. The MFI of phosphorylated proteins at the synapse was quantified. We found a significant decrease of CD3ζ and ZAP-70 phosphorylation signals in INPP5E-KD cells, suggesting that INPP5E sustained initial activation of T cells (Fig 6C and D). Finally, to determine whether impairment of TCR signaling by depleting INPP5E affected effector functions, we measured the secretion amount of IL-2 in the control and INPP5E-KD cells after 24h activation. Consistently, we found that IL-2 secretion in INPP5E-KD cells was significantly attenuated than that in control cells (Fig 6E) (Chan et al., 1995). Thus, these data suggested that INPP5E is essential for early proximal TCR signaling and IL-2 secretion through regulation of CD3ζ and ZAP-70 phosphorylation.

**Figure 6.**
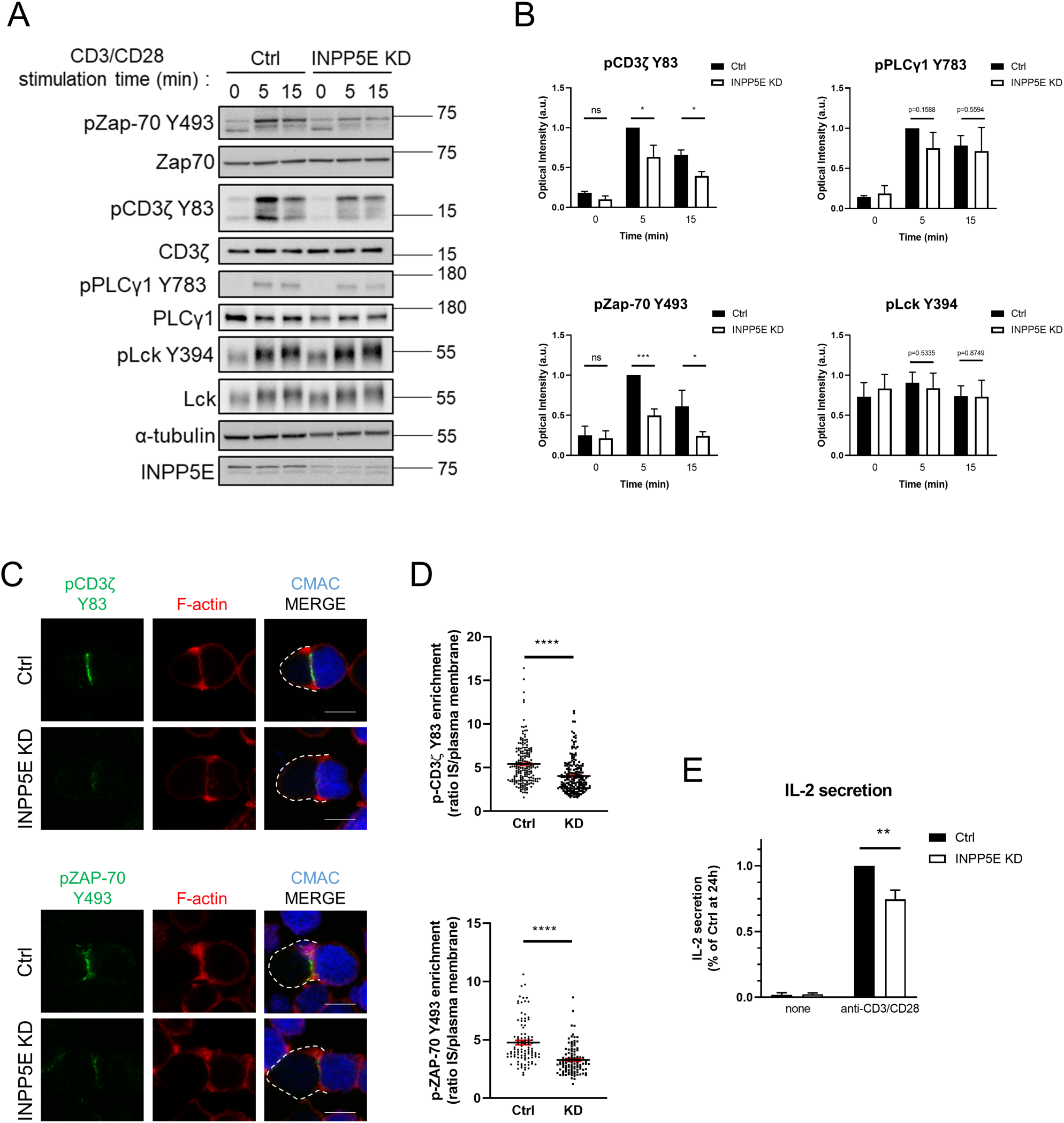
**INPP5E sustains the early proximal TCR signaling and the effector function.** A-B Jurkat cells were transfected with either control (Ctrl) or INPP5E-specific siRNA (KD). Immunoblots showing phosphorylation of indicated proteins (CD3ζ Y83, ZAP-70 Y493, PLCγ1 Y783, Lck Y394) in Ctrl or INPP5E-KD Jurkat T cells activated with crosslinked anti-CD3/CD28 for the indicated times. α-tubulin and non-phosphorylated molecules were blotted as loading controls. Images are representative of three experiments. (B) Quantification of indicated phosphorylated proteins. The results were an average of three independent assays and were all normalized to non-phosphorylated proteins of each sample. Error bars indicate mean ± SD. Paired t-test. ns, not significant. *P < 0.05. C-D Jurkat cells were transfected with either control (Ctrl) or INPP5E-specific siRNA (KD). Immunostaining of pCD3ζ-Y83 (Upper) and pZAP-70-Y493 (Lower) in conjugates of Ctrl or INPP5E-KD Jurkat T cells and CMAC-labeled SEE-pulsed APCs. Cells were co-stained with Alexa Fluor 647 phalloidin to visualize F-actin. Images are representative of three experiments. Quantification of phosphorylated protein enrichment index in conjugates from three independent experiments are showed in (D). n = 183 conjugates for Ctrl, and n = 199 conjugates for INPP5E-KD in pCD3ζ experiments. n = 101 conjugates for Ctrl, and n = 103 conjugates for INPP5E-KD in pZAP-70 experiments. Error bars indicate mean ± SD. Unpaired Student t-test. ****P < 0.0001. E Jurkat cells were transfected with either control (Ctrl) or INPP5E-specific siRNA (KD). ELISA assay showing the amount of IL-2 in the cultured supernatant in Ctrl or INPP5E-KD Jurkat T cells with or without anti-CD3/CD28 for 24 hours. The results were an average of three independent assays and were all normalized to the amount of IL-2 secreted by activated Ctrl cells. *P < 0.05.

## Discussion

Recent studies have demonstrated that some proteins playing roles in the primary cilium are also recruited to the immune synapse (Bustos-Morán et al., 2017; de la Roche et al., 2013), controlling TCR signaling and T cell activation. Here we show that cilium-localized phosphatase INPP5E is indeed enriched at the immune synapse, where INPP5E interacts and regulates CD3 recruitment to the immune synapse by regulating the distribution of PI(4,5)P_2_ and PIP_3_ at the SMAC. Moreover, this regulation can affect the phosphorylation of CD3ζ and ZAP-70 and further enhance IL-2 secretion. Therefore, our results discover that INPP5E is a novel player in the immune synapse, mediating functions by its interplay with TCR complexes.

The dynamics of TCR/CD3 complex at the plasma membrane has been shown to be regulated by phosphoinositides. For instance, PI(4,5)P_2_ level at the surface is significantly lower in Inp54p-overexpressed (referred to as MCID cells) 3A9m T-cells than in wild-type cells. In MCID cells, the lateral diffusion of TCR was significantly augmented, suggesting that PI(4,5)P_2_ dephosphorylation may be required for the movement of the receptors (Chouaki-Benmansour et al., 2018). Moreover, primary T cells that overexpress PIP5K have been shown to possess an increased amount of PI(4,5)P_2_. This consequence thereby enhances T cell rigidity, attenuates and delays the recruitment of TCR complex, and further leads to the impairment of proximal TCR signaling (Sun et al., 2011). The conclusion of these studies is consistent with the results in our works. We observe that PI(4,5)P_2_ level at the center of the synapse was higher in INPP5E-KD Jurkat cells, and the reconstitute CD3ζ-GFP was less recruited to the immune synapse in these cells, suggesting that the diffusion ability of CD3ζ at the plasma membrane was decreased. It remains to be examined whether INPP5E actively regulates PI(4,5)P_2_ or passively maintains the phosphoinositide localization at the synapse.

Modification of phosphoinositides at the plasma membrane also regulates the phosphorylation of proximal TCR components during the formation of the immune synapse (Gagnon et al., 2012; Schamel et al., 2019). CD3ε and CD3ζ contain basic-rich stretches (BRS), allowing these subunits to interact with a series of acidic phospholipids (Bettini et al., 2014; DeFord-Watts et al., 2011, 2009). Mutations in the BRS sequence result in several defects, including impairing the accumulation of CD3 complex and decreasing the TCR-mediated signaling response after antigen engagement. Moreover, BRS in the CD3ε and CD3ζ have been shown to recruit Lck through ionic interactions to initiate CD3 phosphorylation(DeFord-Watts et al., 2009; Li et al., 2017; Zhang et al., 2011). Our studies on INPP5E strongly support the importance of BRS-phosphoinositides interactions in response to T cell activation. INPP5E may prompt CD3ζ phosphorylation via hydrolyzing PI(4,5)P_2_ at the plasma membrane, providing a preferable phosphoinositide microenvironment for the CD3 subunits to be dissociated from the plasma membrane and interact with Lck, and thus promoting the phosphorylation of ZAP-70 and PLCγ1. Notably, INPP5E may not modulate the Lck activity since pTyr 394 level after the anti-CD3/CD28 engagement was not affected when silencing INPP5E in Jurkat cells. As previous research has shown that phosphoinositides at the plasma membrane regulate Lck activities by interacting with its SH2 domain (Sheng et al., 2016), further studies are needed to clarify how INPP5E interplays with CD3ζ and Lck.

Changes of phosphoinositide at the immune synapse appear to be a mirror image of those have been demonstrated in primary cilia (Gawden-Bone et al., 2018; Gawden-Bone and Griffiths, 2019). INPP5E accumulates in the primary cilia, hydrolyzing PI(4,5)P_2_ of the cilia and forming an environment for trafficking and signaling of ciliary proteins (Garcia-Gonzalo et al., 2015; Nozaki et al., 2017; Park et al., 2015; Xu et al., 2017). The immune synapse indeed accommodates some activities similar to those of the primary cilia (Cassioli and Baldari, 2019; Finetti et al., 2011; Griffiths et al., 2010), while results from us and others suggest that there are still different mechanisms involved in these two structures. For example, the TCR recruitment, especially at the early stage, does not require mother centriole polarization and docking. We show that when CD3ζ is recruited to the T cell-APC contact site during antigen engagement, not all the centrioles in T cells are polarized to the immune synapse. While INPP5E is recruited to the immune synapse, centrioles are not necessarily polarized to the cell contact site in some conjugates, suggesting that the accumulation of INPP5E can be independent of the centriole docking. Next, INPP5E ciliary targeting is mediated by a combination of farnesylation and binding to ARL13B, which is at the C-terminal of INPP5E (Fansa et al., 2016; Humbert et al., 2012). Although INPP5E interacts with CD3ζ via its C-terminal portion, however, it utilizes the N-terminal proline-rich domain to target the immune synapse. Together, ciliary machinery seems to be present at the immune synapse, although detailed activities somewhat differ from each other.

In conclusion, we show that INPP5E regulates CD3ζ accumulation by promoting phosphoinositide dephosphorylation at the immune synapse. The changes of tphosphoinositide may prompt the ITAM motifs of CD3ζ to be phosphorylated by Lck and activate ZAP-70. The interplay between INPP5E and CD3ζ facilitates cytokine secretion and T cell activation. Our study unveils how ciliary-enriched phosphatase INPP5E plays an important role in immune synapses and regulates the TCR signaling cascades.

## Materials and Methods

### Cells, plasmids, antibodies and reagents

Jurkat, Clone E6-1 and Raji cells were purchased from BCRC (#60424 and #60116; Bioresource Collection and Research Center, Hsinchu, Taiwan). Cells were maintained in RPMI 1640 (31800-022, Gibco, Thermo Fisher Scientific, Waltham, MA) medium supplemented with 10% FBS (10437-028, Gibco), 2 mM GlutaMAX (35050061, Gibco), and 1% penicillin/ streptomycin (15070063, Gibco). HEK293T cells were cultured in DMEM containing 10% FBS and 1% penicillin/streptomycin.

Jurkat cDNA library were generated by SuperScript IV Reverse Transcriptase (Thermo Fisher), and open reading frames (ORFs) for human CD3ζ and Lck were obtained from this library. CD3ζ was subcloned into pEGFP-N1 vector (Takara Bio, Mountain View, CA), and Lck was subcloned into pEF3 vector, which was constructed by replacing CMV promoter on the pcDNA3 vector into EF1ɑ promoter. The ORFs of these genes were first generated by KOD-Plus-Neo polymerase (TOYOBO, Japan) and then constructed using standard molecular biology techniques. C-terminal Myc-tagged human ZAP-70 was purchased from Sino Biological (HG10116-CY, China). PH-PLCδ-GFP, PH-AKT-GFP, and GFP-PIPKIγ were generous gifts from F.-J. Lee (National Taiwan University, Taiwan), and ORF of PIPKIγ was subcloned into the pmCherry-C1 vector. Tubby-GFP was a kind gift from T. Balla (National Institutes of Health, Bethesda, MD). pSS-FS plasmids, including Flag-tagged INPP5E and the series truncation constructs, were kindly provided by S. Seo (University of Iowa, IA).

The indicated antibodies used for immunoblotting (IB) and immunofluorescence (IF) in this study include: FLAG M2 (F1804; 1:500 IF) and γ-tubulin (T6557; 1:500 IF) from Sigma-Aldrich; pCD3ζ^Y83^ (ab68236; 1:1000 IB, 1:200 IF), and CEP290 (ab84870; 1:200 IF) from Abcam; CD3 (OKT3, 317302, 1:500 IF) from BioLegend; CD3ζ (sc-1239; 1:2000 IB, 1:500 IF), ɑ-tubulin (sc-32293; 1:10000 IB), GM130 (sc-16268; 1:500 IF) and normal rabbit IgG (sc-2027) from Santa Cruz; pZAP-70^Y493^ (#2704; 1:1000 IB, 1:200 IF), PLCγ1 (#5690,1:1000 IB), pPLCγ1^Y783^ (#2821, 1:1000 IB), Lck (#2984, 1:1000 IB), Lck^Y505^ (#2751, 1:1000 IB), and Src^Y416^ (#6943, 1:2000 IB) from Cell Signaling Technology; ZAP-70 (X17371; 1:500 IB) from Genetex; Myc-Tag (AE009; 1:10000 IB) from ABclonal; Centrin (04-1624; 1:500 IF) from Millipore; mCherry (PA5-34974, 1:1000 IF), GFP (A6455; 1:1000 IF, 1:10000 IB), and DYKDDDDK (PA1-984B; 1:2000 IB) from Invitrogen; CEP97 (22050-1-AP; 1:200 IF), CEP164 (22227-1-AP; 1:200 IF), RPGRIP1L (55160-1-AP; 1:200 IF), ARL13B (17711-1-AP; 1:200 IF), and INPP5E (17797-1-AP, 1:200 IF) from Proteintech; and INPP5E (STJ190490, 1:500 IB) from St John’s Laboratory. CEP164 (1F3G10, 1:2000 IF) was kindly provided by C. Morrison (National University of Ireland, Galway).

Staphylococcus enterotoxin E (SEE) was from Toxin Technologies. CellTracker 7-amino-4-chloromethylcoumarin (CMAC, C2110), ProLong Diamond Antifade Mountant (P36965), Alexa Fluor™ 647 Phalloidin (A22287) and highly cross-adsorbed fluorochrome-conjugated secondary antibodies were from Thermo Fisher. Horseradish peroxidase (HRP)-conjugated secondary antibodies were from BioLegend (San Diego, CA). Conformation specific mouse anti-rabbit IgG (#3678) was from Cell Signaling Technology (Danvers, MA). Bovine serum albumin (A9647) and poly-L-lysine (P1274) were from Sigma-Aldrich (St. Louis, MO). Protein G Agarose beads were from Agarose Bead Technologies (Doral, FL). Cy3B maleimide was from GE Healthcare (Pittsburgh, PA).

### Transient transfection

For siRNA knockdown, 2 transfections of 10^6^ Jurkat cells were performed at a 24-h interval using 100 pmol of either control (5’-UUCUCCGAACGUGUCACGU-3’) or INPP5E (5’-GGAAUUAAAAGACGGAUUU-3’; J-020852-05, ON-TARGETplus, GE Dharmacon) siRNA with 100-µl tips from Neon Transfection System (Thermo Fisher) (1,400 V, 10 ms, four pulses). For plasmid transfection, 1 µg of DNA was electroporated in 2.5×10^5^ Jurkat cells 48 h after the first siRNA transfection, using 10-µl tips from Neon Transfection System (1,325 V, 10 ms, three pulses). Cells were harvested and processed for assay 24 h after DNA transfection or 72 h after the first siRNA transfection (Bouchet et al., 2017). For HEK293T, cells were transfected using PolyJet transfection reagents (SignaGen Laboratories, Rockville, MD). 8×10^5^ cells were plated on a 3-cm plate overnight. Cells were transfected with 1 µg plasmid according to the manufacturer’s instructions, and harvested 24 h after transfection. The plasmid pCMV-Tag 4A was used as a Flag tag vector control.

### Cell activation

1×10^6^ (for immunoblot) or 1.8×10^7^ (for immunoprecipitation) Jurkat Cells were incubated with anti-CD3 (1 μg/ml, OKT3, 317325, BioLegend) and anti-CD28 (1 μg/ml, cd28.2, 302902, BioLegend) on ice for 10 min. Antibodies were then crosslinked with donkey anti-mouse IgG (5 μg/ml, 715-005-150, Jackson Immunoresearch, West Grove, PA) on ice for another 10 min. Cells were then activated in 37°C for the indicated times, and the activation was stopped by placing the cells on ice immediately and diluting them with 1 ml cold PBS. Cells were lysed in the lysis buffer supplemented with protease and phosphatase inhibitors, and lysates were centrifuged at 14,000 g for 10 min. Protein concentration was quantified by the Bradford assay (5000006, Bio-Rad, Hercules, CA), and equal amount of proteins were boiled in the SDS sample buffer for immunoblotting.

### Immunofluorescence and image analysis

#### Coverslip preparation

12-mm (0111520) or 18-mm (0111580; Paul Marienfeld, Lauda-Königshofen, Germany) cover glasses were coated with 0.02% poly-L-lysine for 1h at room temperature (RT) and were washed with distilled water. Cover glasses were dried and used in the same day.

#### T cell staining and T cell-APC conjugate preparation

For T cell-APC conjugates of immune synapse experiments, 1×10^7^ cells/ml of Raji cells (used as APCs) were loaded for 1h with 1μg/ml SEE and labeled with 10 μM of the CMAC dye for the last 20 min in 100-μl serum-free RPMI (SFM) at 37°C. Cells were washed with warm PBS for three times and suspended in SFM. When washing Raji cells, Jurkat cells (10^5^ per sample) were allowed to adhere on the coated coverslips for 7 min at 37°C. After that, Raji cells were added to the adhered Jurkat cells for the indicated time. Cells were then fixed in methanol at −20°C for 10 min (for CEP164, CEP97, and ARL13B), or pre-fixed in 2% PFA in PTEM buffer (50 mM Pipes pH 6.8, 25mM HEPES, 10 mM EGTA, 10 mM MgCl2, and 0.1% Triton X-100) at RT for 10 min before methanol fixation (for CEP290, RPGRIP1L, and INPP5E). To detect INPP5E and centriole markers more clearly, adhered Jurkat cells were pre-extracted with PTEM buffer, either with or without Triton X-100, for 15 sec. Cells were then fixed in methanol at −20°C for 10 min. Fixed samples were imaged with epifluorescence microscopy (Olympus X81) with a 100× (NA = 1.30) oil-immersion objective.

For phosphoinositide detection, 2.5 ×10^5^ control, INPP5E-KD, or wild-type Jurkat cells were electroporated with indicated plasmids. After 24h, Jurkat cells (1×10^5^ per 20-μl SFM) were first plated for 7 min at 37°C on 12-mm precoated coverslips. 1×10^5^ SEE-loaded Raji cells (in 20-μl SFM) were then added to each coverslip and co-cultured for 10min. Cells were immediately fixed for 10 min at RT by adding 40-μl 8% PFA, permeabilized with PBST, and blocked as described above. For INPP5E domain mapping, 2.5 ×10^5^ Jurkat cells were electroporated with series truncations 24h before experiments. T cell-APC conjugates were prepared as mentioned in phosphoinositide detection, but with final co-culturing for 20 min. Cells were immediately fixed for 10 min at RT by adding an additional 4% PFA (final 2%), permeabilized with methanol for 20 min, and blocked as described above. For CD3ζ-GFP reconstitution and phosphorylated proteins, 40-μl conjugates (co-cultured for 10min) were fixed for 10 min at RT by adding 40-μl 8% PFA, permeabilized with PBST, and blocked as mentioned above. Confocal images were carried out on a Zeiss LSM 780 using a 100× objective (NA = 1.40). 1 Airy Unit was set as a pinhole for each channel. For Z-series images, sections were set at 0.3-μm intervals.

#### T cell spreading preparation

For spreading T cells on a planar surface, 18-mm precoated coverslips were coated for 2h at 37°C with 1-μg/ml anti-CD3/CD28. Cover glasses were washed three times with warm PBS and keep for 10 min in 37°C. 10^5^ Jurkat cells were then incubated on coverslips for 10 min before being fixed with 4% PFA (Bouchet et al., 2017). Single plane images were acquired on a total internal reflection fluorescence (TIRF) microscope (ELYRA, Zeiss) with a 100× (NA = 1.46) oil-immersion objective lens.

#### Immunostaining

After fixation, cells were washed for 5 min with PBS, permeabilized for 10 min with 0.1% PBST (PBS with 0.05% Triton X-100), and blocked with 3% BSA/PBST for 30 min. Primary antibodies at indicated dilution were prepared with blocking buffer and incubated with cells for 1h. After washing three times in PBST, cells were incubated with Alexa Fluor-labeled secondary antibodies (1:500) and phalloidin dye (1:200) for additional 1h. Finally, cells were washed three times with PBST and mounted with ProLong Diamond Antifade Mountant.

#### Image analysis

ImageJ, Imaris8.2 (Oxford Instruments), and ZEN (Zeiss) software were utilized for image processing and analysis. For control and INPP5E-KD cell comparison, the same acquisition settings and thresholds were used for each image.

For assays related to the recruitment/enrichment of proteins to the synapses at the T cell-APC conjugates, Jurkat cells with similar surface CD3ζ-GFP and Flag-INPP5E levels were chosen for comparing the recruitment ability. For phosphorylated protein enrichment, T cell-APC conjugates from each experiment were randomly selected for processing. Recruitment/enrichment index was measured by a method developed by previous literature (Compeer et al., 2018; Li et al., 2017; Tavano et al., 2004; Zucchetti et al., 2019). Specifically, equal area was selected at the cell-cell contact site (proximal), a region of the T cell opposite from the cell contact site (distal), and a background area outside of the cell (BG). The recruitment index was calculated with the formula: [MFI at the proximal site – BG] / [MFI at the distal site − BG]. Quantification of MFI was performed with Image J. For recruitment of INPP5E truncation, we scored each conjugate as positive when most of the Flag signals were accumulated at the cell contact.

For recruitment of endogenous INPP5E to the pseudo-synapse on TIRF images, spreading cells were chosen randomly from each experiment, and the area for measurement was selected by the contour of CD3ζ. Image J was used to quantify the MFI.

For the quantification of phosphoinositide clearance, Z-stack confocal images were reconstructed into 3D images by Imaris software, and the en face view at the contact site was used for analysis. Clearance ability was accessed following previous studies (Jenkins et al., 2014; Schubert et al., 2012). Briefly, ‘cleared’ was scored when a continuous GFP ring was formed at the center of the synapses; any interruption at the center was classified ‘not cleared’.

#### dSTORM imaging and image analysis

Super-resolution dSTORM imaging was performed as previously described (Chong et al., 2020; Yang et al., 2018). Briefly, Alexa-Fluor 647 labeled anti-INPP5E (1:200) and Cy3B-conjugated anti-CD3ζ (1:100) were used for immunostaining and imaged with a modified inverted microscope (Eclipse Ti-E, Nikon, Tokyo, Japan). The dSTORM imaging buffer included Tris-NaCl buffer at pH 8.0, and an oxygen-scavenging system consisting of 10 mM mercaptoethylamine (MEA, 30070, Sigma-Aldrich) at pH 8.0, 0.5 mg/mL glucose oxidase (G2133, Sigma-Aldrich), 40 mg/mL catalase (C40, Sigma-Aldrich), and 10% (w/v) glucose (G8270, Sigma-Aldrich). Fiducial markers (0.1 μm Tetraspeck, Thermo Fisher) were used to correct the drift. Typically, 15,000 frames were recorded every 20 ms (exposure time). Individual single-molecule peaks were then real-time localized using a MetaMorph Super-resolution Module (Molecular Devices), based on a wavelet segmentation algorithm. Super-resolution images in figures were cleaned with a Gaussian filter with a radius of 1 pixel. Alexa 647 channel was first recorded, and then the Cy3B channel was acquired with the corresponding filter (593/40, Chroma).

### Immunoblotting

Cells were washed with PBS twice and lysed in the ice-cold lysis buffer (20 mM Tris-HCl, pH7.5, 150 mM NaCl, 1 mM EDTA, 1% NP-40) that contains protease inhibitors and PhosSTOP inhibitors (Sigma-Aldrich). Cells were then centrifuged at 14,000 g for 10 min at 4°C to remove cell debris. Protein concentrations were determined by the Bradford protein assay (Bio-Rad). Equal amounts of proteins were mixed with the SDS sample buffer, boiled at 95°C for 5 min, and resolved by SDS-PAGE. The separated proteins were then transferred to the PVDF membranes (0.45 µm Immobilon®-E; Merck Millipore). Membranes were blocked either with 2.5% nonfat milk in TBST (50 mM Tris, pH 7.5, 150 mM NaCl, and 0.05% Tween-20) or 3% BSA (for phospho antibodies) for 1h at RT and incubated with primary antibodies overnight at 4°C. Blots were washed three times with TBST and incubated with HRP-conjugated secondary antibodies (Biolegend) for 1h at RT. After washing three times with TBST, proteins were visualized with Western Lightning Plus substrate (PerkinElmer).

### Immunoprecipitation

For immunoprecipitation of endogenous INPP5E in Jurkat cells, cells were first activated as described in the methods above. After activation, cells were immediately mixed with ice-cold 5x lysis buffer supplied with protease inhibitors. Cells were centrifuged at 14,000 g for 10 min at 4°C to remove cell debris. Cell lysates were then pre-cleared with 20-μl protein G agarose beads for 1h to remove antibodies for activation. 800-μg lysates were suspended in 400-μl lysis buffer and immunoprecipitated with 1-μg anti–INPP5E antibody (Proteintech, Rosemont, IL) overnight, and incubated with 20-μl protein G agarose beads for additional 1h on the next day. The beads were washed three times with the lysis buffer, eluted by boiling in 2x sample buffer, and resolved by SDS-PAGE.

For immunoprecipitation in HEK293 cells, 8×10^5^ cells were transfected as described above. After 24-h transfection, cells were washed two times with ice-cold PBS, and lysed in the ice-cold lysis buffer. 500-μg of pre-cleared lysates were immunoprecipitated with 1-μg anti-FLAG antibody (Sigma-Aldrich) overnight, and incubated with 20-μl protein G agarose beads for another 1 h. The beads were washed three times with the lysis buffer with 0.01% glycerol, eluted by sample buffer and resolved by SDS-PAGE.

### ELISA

1×10^5^ Jurkat cells were stimulated with soluble anti-CD3/CD28 (each concentration of 1 μg/ml) in 96-well plates for 24 h, and supernatants were collected for the ELISA assay. IL-2 was quantitated with human IL-2 ELISA Set (OptEIA, 555190, BD bioscience) following the manufacturer’s protocols.

### Statistical Analysis

Unpaired Student’s t-test (parametric) and p-value was performed with GraphPad Prism 8.0 software. Error bars indicate SDs unless otherwise specified.

## Acknowledgments

We thank Dr. Jen Liou (The University of Texas Southwestern Medical Center, Dallas, TX) for discussion and critical comments. We thank Sue-Ping Lee (Imaging Core at Institute of Molecular Biology, Academia Sinica) for assisting with technical assistance of TIRF microscopy. We also thank the staffs of Technology Commons (College of Life Science, National Taiwan University) for help with confocal microscopy. This work was supported by the Ministry of Science and Technology, Taiwan (MOST 107-2313-B-001-009, 108-2313-B-001-003) and National Taiwan University and Academia Sinica Innovative Joint Program Grant (NTU-SINICA-108L104303) to TYC,FHY, WMC, and JCL; by the National Health Research Institutes (NHRI-EX109-10610BC) and National Taiwan University and Academia Sinica Innovative Joint Program (109L104303) to HJT; by the MOST (109-2628-B-010-015, 109-2313-B-010-001) to WJW.

## Author Contributions

Author contributions: TYC, WJW, and JCL designed research; TYC performed research; TYC, FHY, and WMC analyzed data; and TYC, HCT, WJW, and JCL wrote the paper.

## Conflict of Interests

The authors have declared that no conflict of interest exists.

## Supplementary results

**Figure S1.**
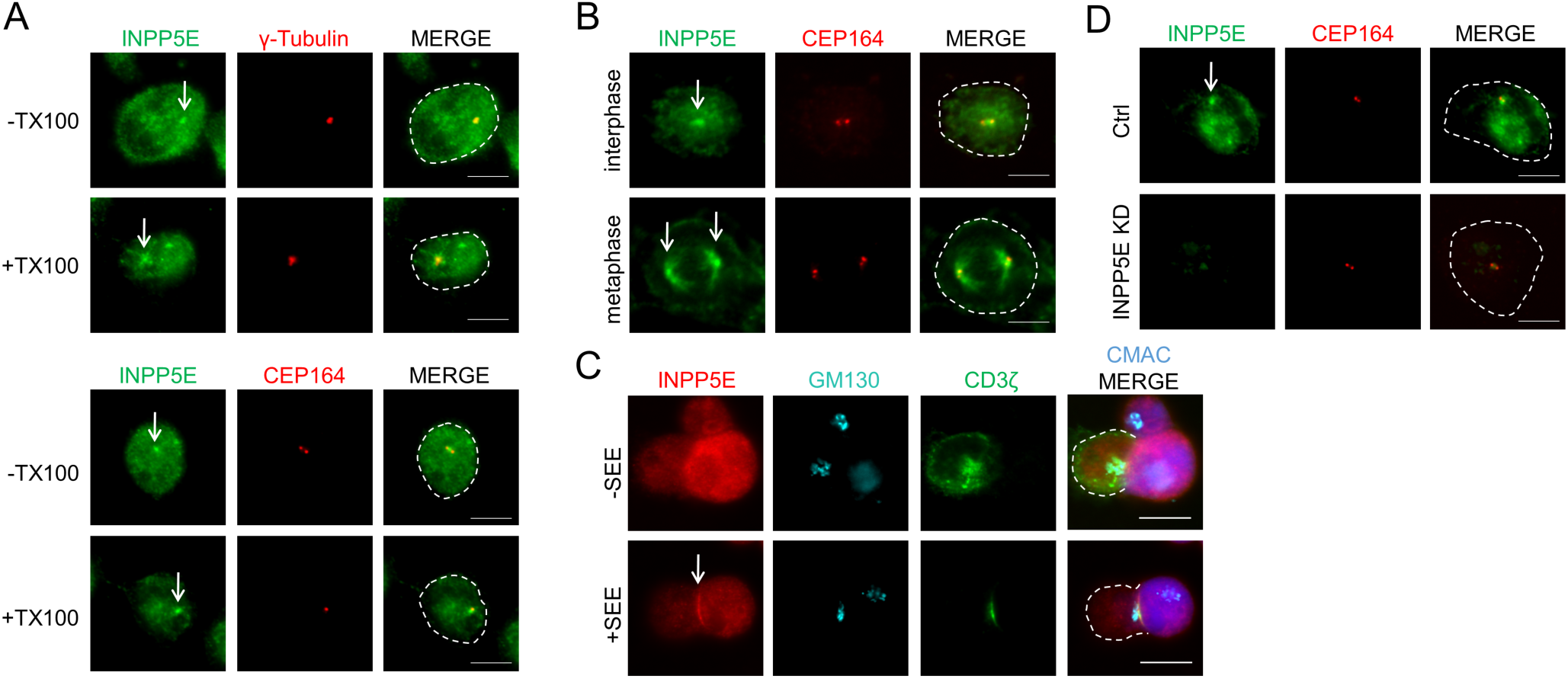
A Immunostaining of INPP5E in Jurkat cells, either without or with Triton X-100 extraction before fixation. Cells were co-stained with anti-γ-tubulin or anti-CEP164 antibodies as centriole markers. B Immunostaining of INPP5E in Jurkat cells in prometaphase (Upper) and anaphase (Lower). C Immunostaining of INPP5E in Jurkat cells. Cells were co-stained with the anti-GM130 antibody as the Golgi marker. D Immunostaining of INPP5E in scramble control or INPP5E-specific siRNA knockdown Jurkat cells. Cells were co-stained with anti-CEP164 antibodies. Scale bar = 5 μm.

**Figure S2.**
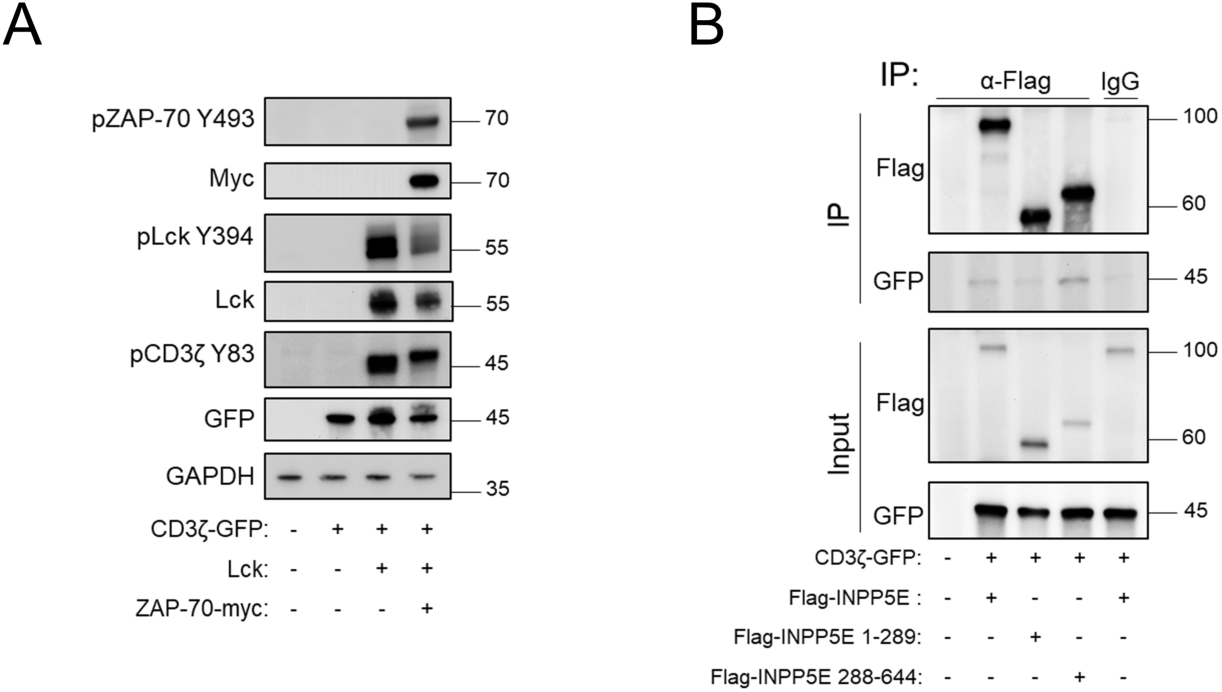
A HEK293T cells were transfected with listed constructs. Cell lysates were resolved by SDS-PAGE and immunoblotted with specific antibodies. B HEK293T cells were co-expressed with CD3ζ-GFP and INPP5E deletion mutant. Cells were then lysed and immunoprecipitated (IP) with anti-Flag antibody. The immuno-precipitates were resolved by SDS-PAGE and immunoblotted with indicated antibodies. Images are representative of at least three experiments.

**Figure S3.**
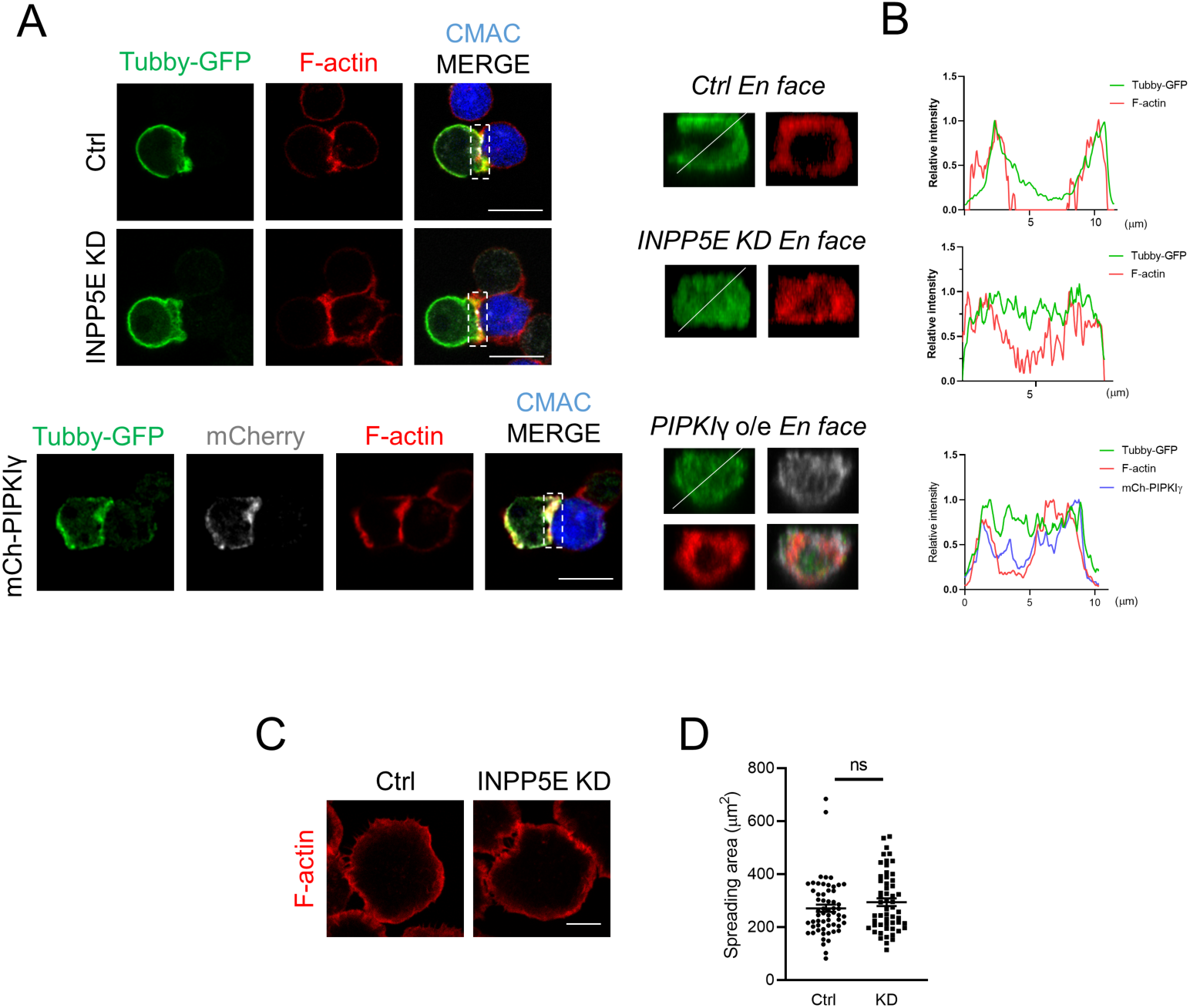
A-B Immunostaining of Tubby-GFP in conjugates of Jurkat T cells and CMAC-labeled SEE-pulsed APCs. (A) Images are shown in the xy plane (scale bar, 10 μm) or 1.0 μm 3D-reconstructions en face at the T cell-APC contact site in the xz plane (scale bar, 3 μm). Cells were stained with anti-GFP antibody. Alexa Fluor 647 phalloidin was stained to visualize F-actin. Dotted white squares indicate the regions that were selected for 3D reconstruction. Upper, control siRNA transfected cells; middle, INPP5E-specific siRNA transfected cells; lower, mCherry-PIPKIγ overexpressed (o/e) cells. Images are representative of two experiments. (B) Line profiling (derived from the white lines in the merged en face images) showing the fluorescence intensities of F-actin, Tubby-GFP, and mCherry-PIPKIγ. C-D Jurkat cells were transfected with either control (Ctrl) or INPP5E-specific siRNA (KD). Immunostaining of F-actin in spreading Jurkat cells on anti-CD3/ CD28-coated coverslips for 10 min, and measurement of spreading area (D) from three independent experiments. n = 59 cells for control, and n = 55 cells for INPP5E-knockdown. Scale bar = 5 μm. Error bars indicate mean ± SD. Unpaired Student t-test. Images are representative of three experiments.

